# Lattice defects induce microtubule self-renewal

**DOI:** 10.1101/249144

**Authors:** Laura Schaedel, Denis Chrétien, Charlotte Aumeier, Jérémie Gaillard, Laurent Blanchoin, Manuel Théry, Karin John

## Abstract

The dynamic instability of microtubules is powered by the addition and removal of tubulin dimers at the ends of the microtubule. Apart from the end, the microtubule shaft is not considered to be dynamic. However recent evidence suggests that free dimers can be incorporated into the shaft of a microtubule damaged by mechanical stress. Here we explored whether dimer exchange was a core property of the microtubule lattice independently of any external constraint. We found that dimers can be removed from and incorporated into the lattice at sites along the microtubule shaft. Furthermore, we showed by experiment and by modeling that rapid dimer renewal requires structural defects in the lattice, which occur in fast growing microtubules. Hence long-lived microtubules have the capacity to self-renew despite their apparent stability and thereby can potentially regulate signaling pathways and structural rearrangements associated with tubulin-dimer exchange at sites along their entire length.

## INTRODUCTION

The discovery that microtubules have the property of dynamic instability came from the investigation of microtubule-growth kinetics in response to the dilution of soluble tubulin (Mitchison and Kirschner, 1984). Dynamic instability allows microtubules to co-exist in the growing and shrinking states and to abruptly transit between the two. Concurrent with this discovery, an intense controversy animated the investigation of the location of tubulin incorporation into microtubules (Inoue, 1981), which was proposed to occur along lateral sides (Salmon et al., 1984) or at the end of microtubules (Leslie et al., 1984). Soon after, the injection of labeled tubulin dimers in living cells revealed that microtubule growth was mainly promoted by dimer addition at the microtubule ends (Soltys and Borisy, 1985). Further experiments showed that GTP hydrolysis in those newly added dimers at the growing end, triggered the transition to the shrinking phase (Duellberg et al., 2016). Hence microtubule dynamics appeared to be fully controlled by tubulin exchange at microtubule free ends (Akhmanova and Steinmetz, 2008). However, this unequivocal scenario was contested by the observation that end-stabilized microtubule subjected to free tubulin washout fold and broke along the shaft of the microtubule, reviving the possibility of lateral exchange of dimers at these sites (Dye et al., 1992). This hypothesis was recently demonstrated by experiments on microtubules subjected to repeated cycles of bending. Microtubules appeared capable of self-repair and maintaining rigidity over repeated cycles by incorporating free tubulin dimers into sites of damaged lattice (Schaedel et al., 2015). Later, live cell imaging revealed that those damaged sites in which tubulin dimers were incorporated also act as rescue sites protecting the microtubule from further disassembly and promoting the resumption of growth (Aumeier et al., 2016; de Forges et al., 2016). The hypothesis of lateral exchange of tubulin dimers along the microtubule shaft had been predicated on the notion that the lattice bending and breakage would trigger the exchange. However, lattice self-repair has been observed under rather small hydrodynamic drag forces (1 pN on 30 μm long microtubules, corresponding to an average work of about 2 *k*_*B*_*T* performed on a dimer) (Schaedel et al., 2015, and Supplementary Material). This suggests that such dimer exchange can occur at a slower time scale in response to thermal stochastic forces only. This modified hypothesis would then apply to almost all long-lived cellular microtubules.

## RESULTS

We tested the possibility of spontaneous microtubule turnover by growing microtubules *in vitro* from purified brain tubulin (Weisenberg, 1972). Dynamic microtubules were nucleated from short taxol-stabilized microtubule seeds and elongated in the presence of 20 μm of free tubulin dimers (“*tubulin growth” condition*), 10% of which was labeled with red fluorophores (Fig. 1a, step I). Microtubule seeds were grafted on micropatterned sites surrounded by anti-fouling coating to maintain microtubule end anchorage to the substrate, and thus preventing microtubule displacement when switching medium, and avoiding microtubule shaft interaction with the substrate (Portran et al., 2013; Schaedel et al., 2015). Importantly, and in contrast with common practices in the field, microtubule shafts were not stabilized with taxol because it can impact lattice structure (Kellogg et al., 2017; Yajima et al., 2012) and because its stabilizing effect can impact intrinsic biochemical properties of the lattice (Reid et al., 2017). Microtubule ends where then exposed to free tubulin in the presence of GMPCPP, a non-hydrolysable analog of GTP to protect them from depolymerization (Fig 1a, step II). Microtubule ends were capped to enable the investigation of GDP lattice turnover over a longer period than the typical lifetime of microtubules governed by dynamic instability in vitro. Capped microtubules were then exposed to 20 μM of free tubulin dimers labeled with a green fluorophore for 15 minutes (“*tubulin exchange*”*condition*) (Fig 1a, step III). After washout of the free tubulin (Fig 1a, step IV), green tubulin spots became apparent along microtubule shafts (Fig. 1b). This method revealed that tubulin incorporation can occur spontaneously, ie in the absence of external bending forces, in less than 15 minutes.

**Figure 1:**
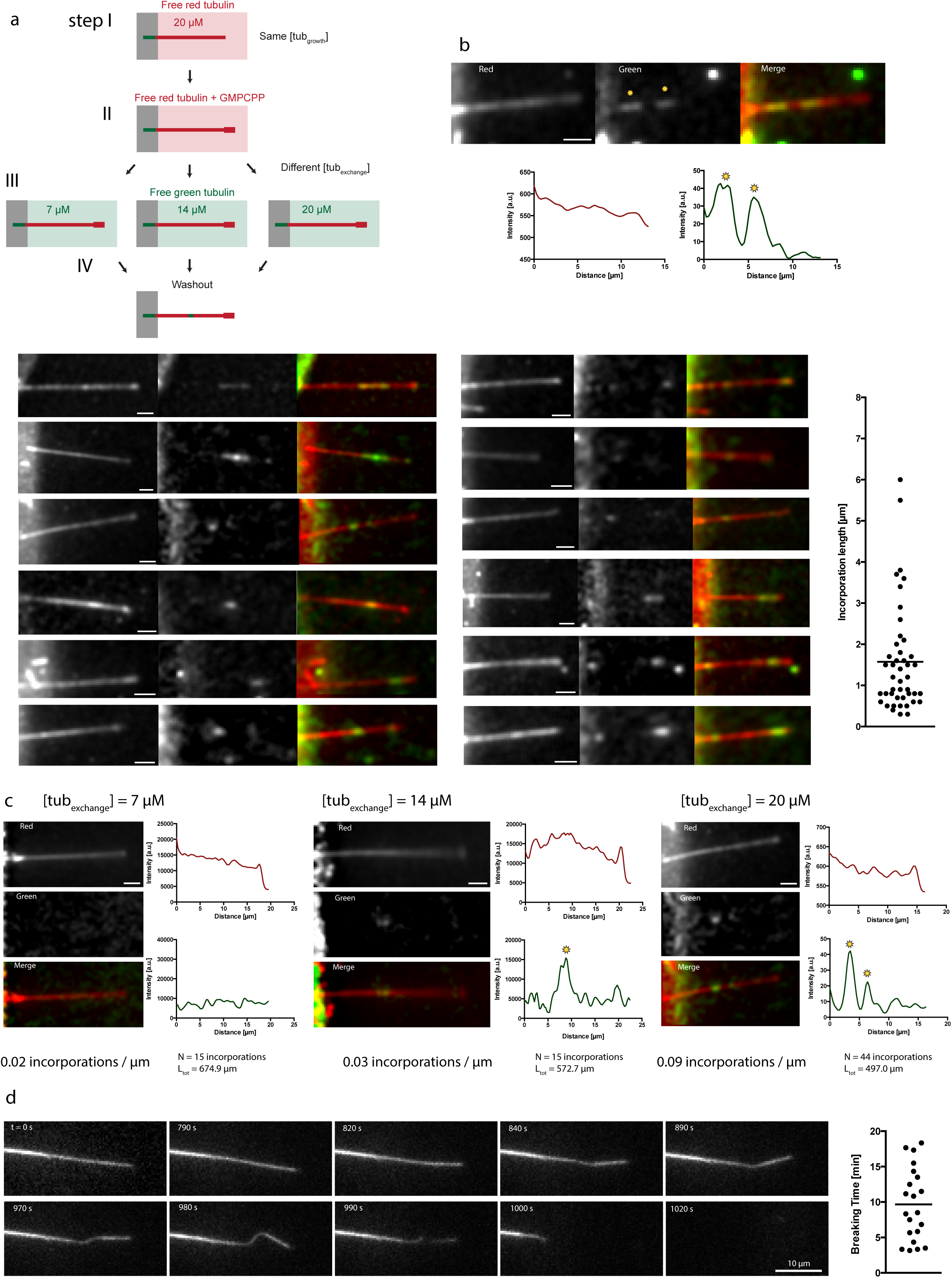
(a) The schematic representation of the experimental setup used to test microtubule-lattice turnover in the absence of external forces. Microtubules were grown with red-fluorescent tubulin at a concentration of 20 μM before they were capped with GMPCPP and exposed to varying concentrations of green-fluorescent free tubulin. After 15 min, the green tubulin was washed out to reveal spots of green tubulin along the red microtubule. (b) Examples of red microtubules showing spots of incorporated green tubulin along the lattice. The graphs represent line scans along the microtubule (red curve: red fluorescent channel, green curve: green fluorescent channel). Sites of incorporated tubulin were identified as zones of fluorescence intensity, which were at least 2.5-fold higher than the background fluorescence (see yellow stars). The graph on the right shows the length of the incorporated tubulin stretches. The bar represents the mean length. Scale bars: 3 μm. (c) Microtubules grown at 20 μM tubulin concentration were exposed to 7 μM, 14 μM and 20 μM of green-fluorescent free tubulin. The images and line scans represent typical examples for every concentration. The average frequency of incorporation spots is indicated below. Scale bars: 3 μm. (d) Sequence of images showing a capped microtubule in the absence of free tubulin, which develops a soft region that extends along the microtubule before breakage. The graph on the right shows the time that microtubules survive before breaking, the bar represents the mean survival time.

Tubulin dimer incorporation appeared to be a cooperative process because the sites of incorporation ranged in length from less than 1 μm up to 6 μm, which suggested up to several hundreds of dimers were included (Fig. 1b). For dynamic microtubules, the law of mass action requires that the polymerization speed at the growing microtubule end depends on the concentration of free tubulin (Mitchison and Kirschner, 1984). We therefore hypothesized that tubulin incorporation into the lattice would also depend on the concentration of free tubulin. To test this hypothesis, microtubules generated in the *tubulin-growth* process were exposed to different concentrations of free-tubulin in the *tubulin-exchange* process. Sites of incorporated tubulin were identified using line scans of green-fluorescence intensity along the microtubule and corresponded to zones of fluoresecence intensity that were at least 2.5-fold higher than the background fluorescence. At subcritical (7 μM) and critical concentrations (14 μM), the frequencies of tubulin-incorporation sites were 0.02/μm and 0.03/μm, respectively. At 20 μM, the frequency increased to 0.09/μm (see Fig. 1c). Therefore, tubulin incorporation into the shaft is dependent on the concentration of free tubulin, as has been found at end of the microtubule.

A genuine turnover process would imply that some dimers also leave the lattice. This means that, if no incorporation occurs, microtubules should exhibit marked dimer loss. To verify this, we monitored end-capped microtubules in the absence of free tubulin in the solution. We found that microtubules developed soft, flexible regions a few micrometers in length, which continued to spread along the microtubule shafts. The highly bent regions eventually broke after 10 ± 5 min, followed by rapid shortening of the remaining microtubule ends (Fig. 1d). This indicated that tubulin is lost preferentially in a longitudinal direction and in a cooperative process, as expected for an anisotropic lattice where longitudinal bonds are stronger than lateral bonds (Sept et al., 2009).

The incorporation and loss of tubulin in microtubules in the absence of external constraints suggested a genuine spontaneous and continuous process of tubulin turnover that depended on the concentration of free tubulin, similar to the gain and loss of tubulin at microtubule ends. However, previous work has suggested that the lattice energy is too high to permit a spontaneous turnover (VanBuren et al., 2002; VanBuren et al., 2005; Sept et al., 2009). We re-investigated this question by performing Monte-Carlo simulations of the canonical microtubule lattice (13 protofilaments and 3 start left-handed helix) (Mandelkow et al., 1986; Chrétien and Wade, 1991) in a comparable approach to what has previously been modelled (VanBuren et al., 2002; Wu et al., 2009; Gardner et al. 2011 and Methods). However, we specifically allowed for the exchange of dimers between the microtubule shaft and the surrounding solution, a possibility which was not considered previously. Also, exchange depended critically on the presence of nearest neighbors in the lattice, via Arrhenius type kinetics (Fig. 2a and Methods). The parameters for the modelling were adapted from Gardner et al. (Gardner et al., 2011) to approximately match experimentally measured polymerization and depolymerization speeds, such that microtubules exhibited a dynamic instability for tubulin concentrations close to the critical concentration (see Fig. 2b, Table S1 and Supplementary Fig. S1). To model the experiments of end-stabilized microtubules (Fig. 1b), we simulated the exchange of dimers between 20 μm long microtubule shafts and the solution containing 20 μM free tubulin. The frequency of dimer exchange between the lattice and pool of free dimers during the first 15 minutes was considered too low to be detectable experimentally (Fig. 2c). The simulation showed an exchange of about 1 dimer per helical turn would rather take several hours and be distributed nearly homogeneously along the microtubule (Fig. 2d). We concluded that this model of a regular lattice, where all dimers interact with four neighbors, was not sufficient to explain the localized exchange patterns.

**Figure 2:**
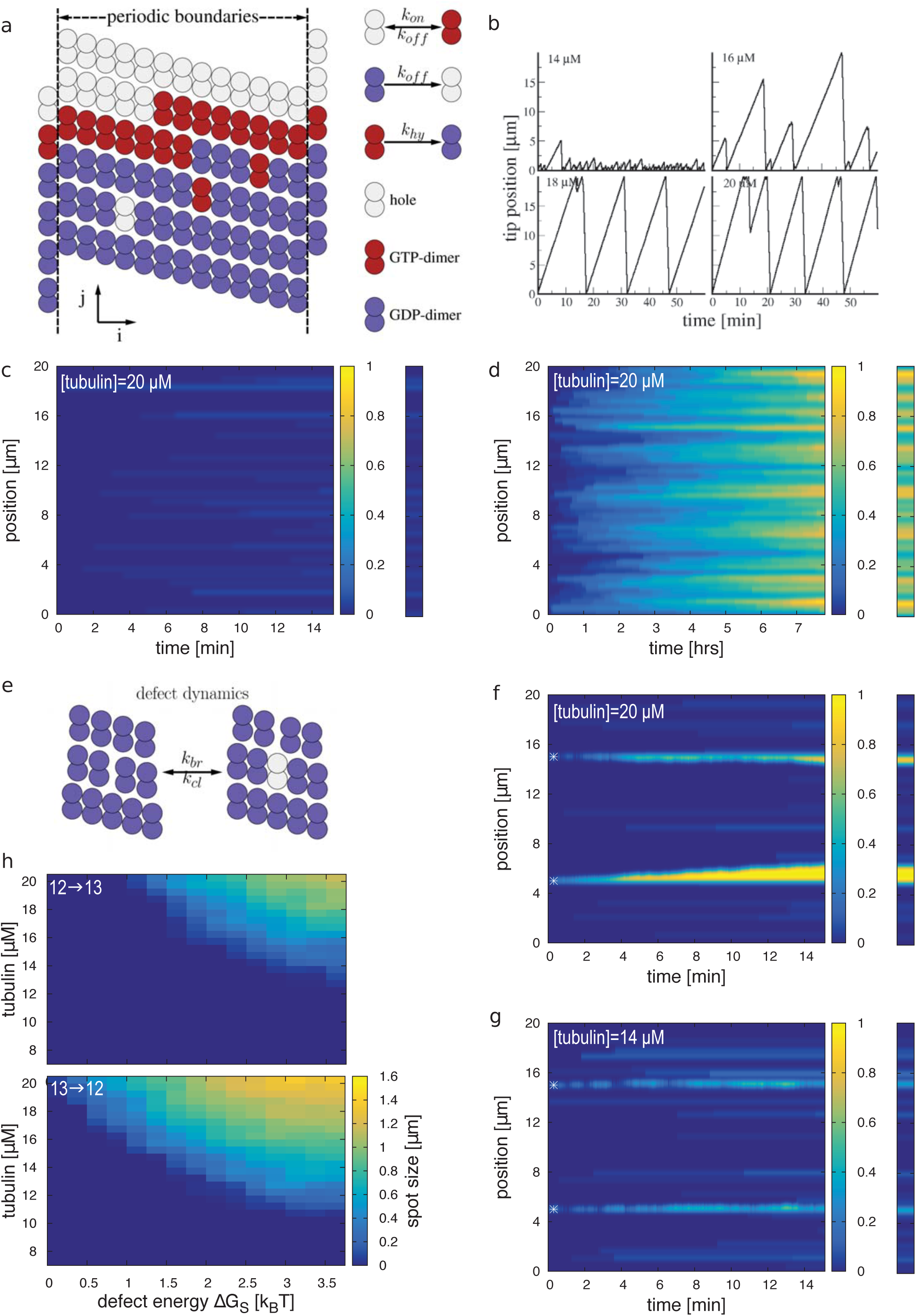
Monte-Carlo simulations of microtubule lattice dynamics. (a) Lattice scheme for a 13_3 protofilament lattice with a seam and possible lattice transitions. (b) Tip dynamics for various concentrations of free tubulin as indicated in the legends. (c,d) Exchange of tubulin dimers between the microtubule shaft of end-stabilized 13_3 microtubules and the surrounding medium with 20 μM free tubulin at short (c) and long (d) time scales. (e) Scheme of the breathing mechanism of the microtubule lattice at a dislocation defect. (f,g) Exchange of tubulin dimers between the microtubule shaft and the surrounding medium with 20 μM (f) and 14 μM (g) free tubulin at sites of dislocation defects. The initial position of the defect is marked by a white star. *ΔG*_*S*_=2 *k*_*B*_*T*. The bar on the right of (c,d,f,g) shows at snapshot of the exchange pattern after 15 min as it would be observed experimentally. The color code of (c,d,f,g) indicates the number of exchanged dimers per helical turn. (h) Observable size of incorporation spots in μm of 12⇒13 (top) and 13⇒12 transitions after 15 min as a function of the free tubulin concentration and the defect strain energy *ΔG*_S_.

The experimentally observed incorporation patterns, made of few long stretches rather than numerous dispersed spots (Figs. 1b,c), suggested that some lattice sites might be structurally different from other parts of the lattice and favor the local exchange of tubulin. As a matter of fact, the microtubule lattice is not perfect but exhibits defects (Chrétien and Fuller, 2000; Chrétien et al., 1992; Vitre et al., 2008; Doodhi et al., 2016; Schaap et al., 2004), such as missing dimers, lateral cracks and dislocation defects. The most common defect is the dislocation defect, where the protofilament number and/or the helix start number change along the shaft of the microtubule and which are typically separated by several micrometers (Chrétien et al., 1992). We hypothesized that the elastic stress, associated with dislocation defects, might be sufficient to promote spontaneous tubulin turnover at that site in the microtubule shaft and to explain the localized exchange pattern. We therefore extended our Monte-Carlo model to simulate the microtubule dynamics at transitions between 12 and 13 protofilament lattice conformations via dislocation defects. Thereby we postulated (i) that a passive lattice breathing mechanism occurs at the defect site (Fig. 2e); (ii) that there is an elastic strain energy *ΔG*_*S*_ associated with the defect; and (iii) that the 13 protofilament lattice is slightly more stable than the 12 protofilament lattice (Hunyadi et al., 2005, see also Methods). We simulated the exchange of dimers for microtubule shafts containing two dislocation defects, a 13⇒12 and a 12⇒13 transition (in the direction of the microtubule plus end). For a high free tubulin concentration (Fig. 2f), we found that free protofilaments at the dislocation site elongated by dimer addition, albeit in the sense that free plus ends were elongating faster than free minus ends. Elongation occured with a speed of 1-2 μm within 15 min. At a low tubulin concentration (Fig. 2g), the elongation of the free protofilament ends was below the detection limit. Growth speed of plus and minus protofilament ends at the dislocation defect was dependent on the dislocation energy penalty *ΔG*_*s*_ and the free tubulin concentration (Fig. 2h). Overall, rapid tubulin turnover at the defect was a robust behavior, which can be simulated over a broad range of parameters in the Monte Carlo model (see Supplementary Fig. S1). Hence the numerical simulations show that small lattice imperfections with little impact on overall microtubule appearance (width and curvature) are sufficient to promote turnover. They also predict that the frequency of dislocation defects should directly impact the frequency of the sites of tubulin turnover.

Lattice defects have been previously observed by cryo-electron microscopy on microtubules assembled from purified tubulin (Chrétien *et al.*, 1992; Chrétien and Fuller, 2000). Here, we investigated the effect of tubulin concentration, and hence of microtubule growth rate, on the frequency of lattice defects (Fig. 3). Microtubules were nucleated from centrosomes to facilitate visualization of their elongating plus ends. They were assembled at three tubulin concentrations (6.5 μM, 13 μM and 19.5 μM) and were vitrified at different times during the assembly process. We could characterize the growth rate and the lattice-defect frequency as a function of tubulin concentration after 3 min of assembly (Fig. 3a). Transitions types were preferentially 13⇒12, 13⇒14 and 14⇒13, although transitions that involved the loss of two protofilaments or only a change in the helix-start number were also observed (Supplementary Table S2). At the lowest tubulin concentration the microtubule growth rate was 0.8 ± 0.4 μm.min^-1^ and the defect frequency was about 0.006 μm^-1^ (Fig. 3a and supplementary Table S2). At a two‐ and threefold higher tubulin concentration, microtubules grew at 2.2 ± 0.9 μm.min^-1^ and 3.5 ± 0.9 μm.min^-1^ respectively, and the dislocation frequency increased to 0.014 μm^-1^ and 0.45 μm^-1^, respectively. These results demonstrated that the lattice-defect frequency increased with increased free-tubulin concentration, and hence with increased microtubule growth rate.

**Figure 3:**
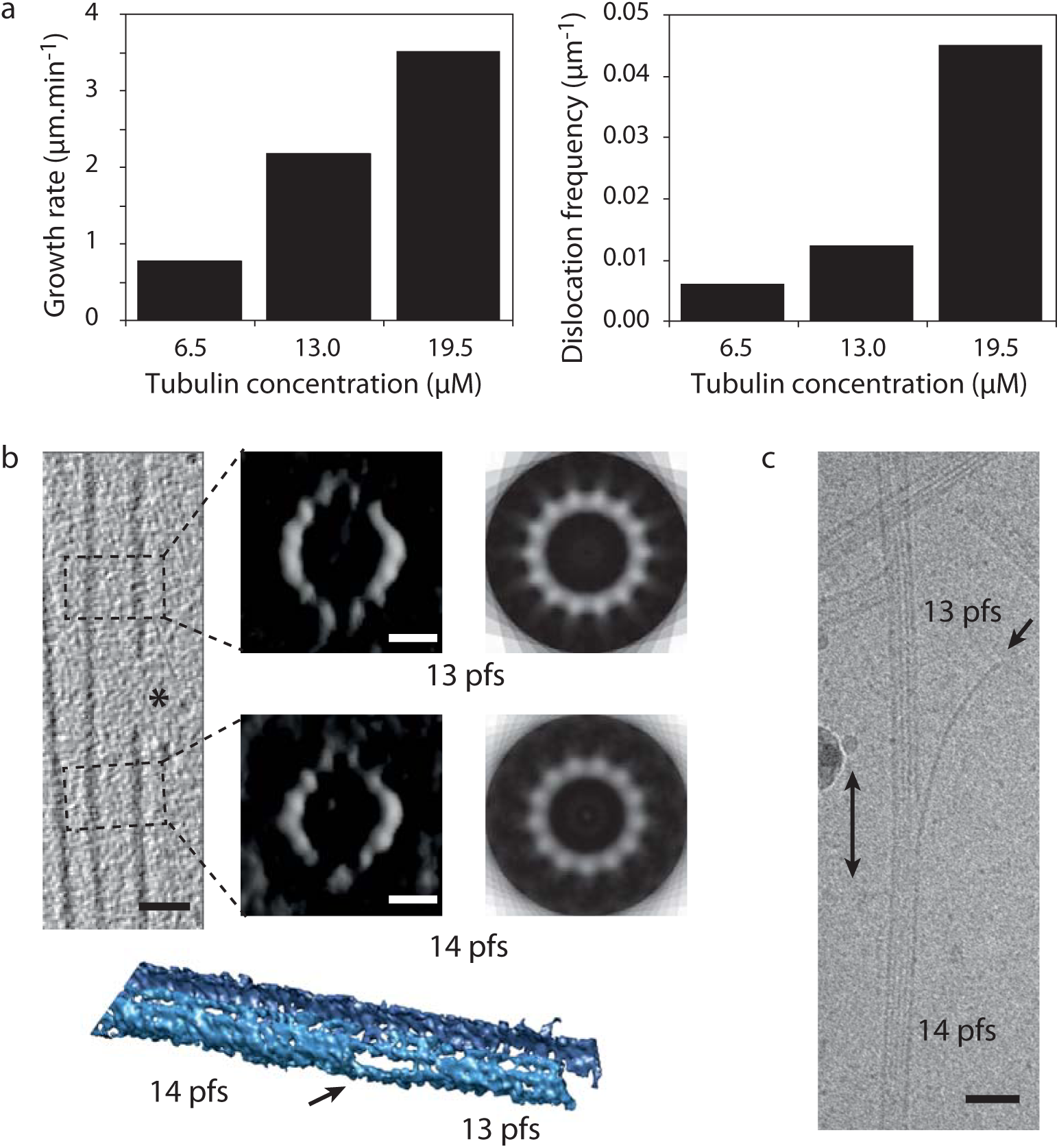
(a) Growth rate (left) and dislocation frequency after 3 min of assembly (right) in centrosome-nucleated microtubules as a function of the free tubulin concentration. (b) Cryoelectron tomogram of a 13 ⇔14 protofilament-number transition. Longitudinal slice through the tomogram at the transition region (top left). Scale bar: 25 nm. The star indicates the open portion of the microtubule wall. Transverse slices (top middle) and rotational averaging of these images (top right) before and after the transition (boxed regions in the left image). Scale bars: 10 nm. 3D rendering of the dislocation (bottom) visualized under UCSF Chimera (Pettersen *et al.*, 2004). The arrow points to the dislocation. pfs: protofilaments. (c) Outwardly curved protofilament extension (arrow) at a 13 ⇔14 protofilament number transition (double arrow). Scale bar: 50 nm.

Changes in protofilament number necessitate the opening of the microtubule lattice. This can be visualized in three dimensions (3D) based on microtubule reconstructions from cryo-electron tomography at medium resolution and in the absence of averaging methods (Coquelle *et al.*, 2011). With this approach, the characteristic transition from 13 to 14 protofilaments can be observed (Fig. 3b), with the transition region marked as a ~75 nm length gap in the microtubule lattice (Fig.3b). Also, small protofilament extensions can occasionally be observed in the transition region (Fig. 3c, and Chrétien and Fuller, 2000). The structure of these extensions is similar to that of the tubulin sheets observed at growing microtubule ends (Chrétien *et al.*, 1995; Guesdon *et al.*, 2016). These observations suggest that tubulin can indeed be added at the end of free protofilaments at the transition regions.

We then took advantage of the relationship between microtubule growth rate and lattice-defect frequency to modulate the latter and test its direct contribution to the regulation of tubulin turnover along the microtubule shaft. Thus, red microtubules were grown in the presence 14 μM, 20 μM and 26 μM of free tubulin, capped with GMPCPP (Fig. 4a, steps I and II) and exposed to the same concentration of green tubulin (20 μM) for 15 min (step III) before washout of the free tubulin (step IV). Sites of incorporated tubulin were quantified: microtubules grown at 14, 20 and 26 μM had on average, 0.05, 0.09 and 0.17 sites of tubulin incorporation per micrometer, respectively (Fig. 4b), the latter corresponding to 1 incorporation site every 6 μm. This demonstrated that higher defect frequencies were associated with increased tubulin turnover.

**Figure 4:**
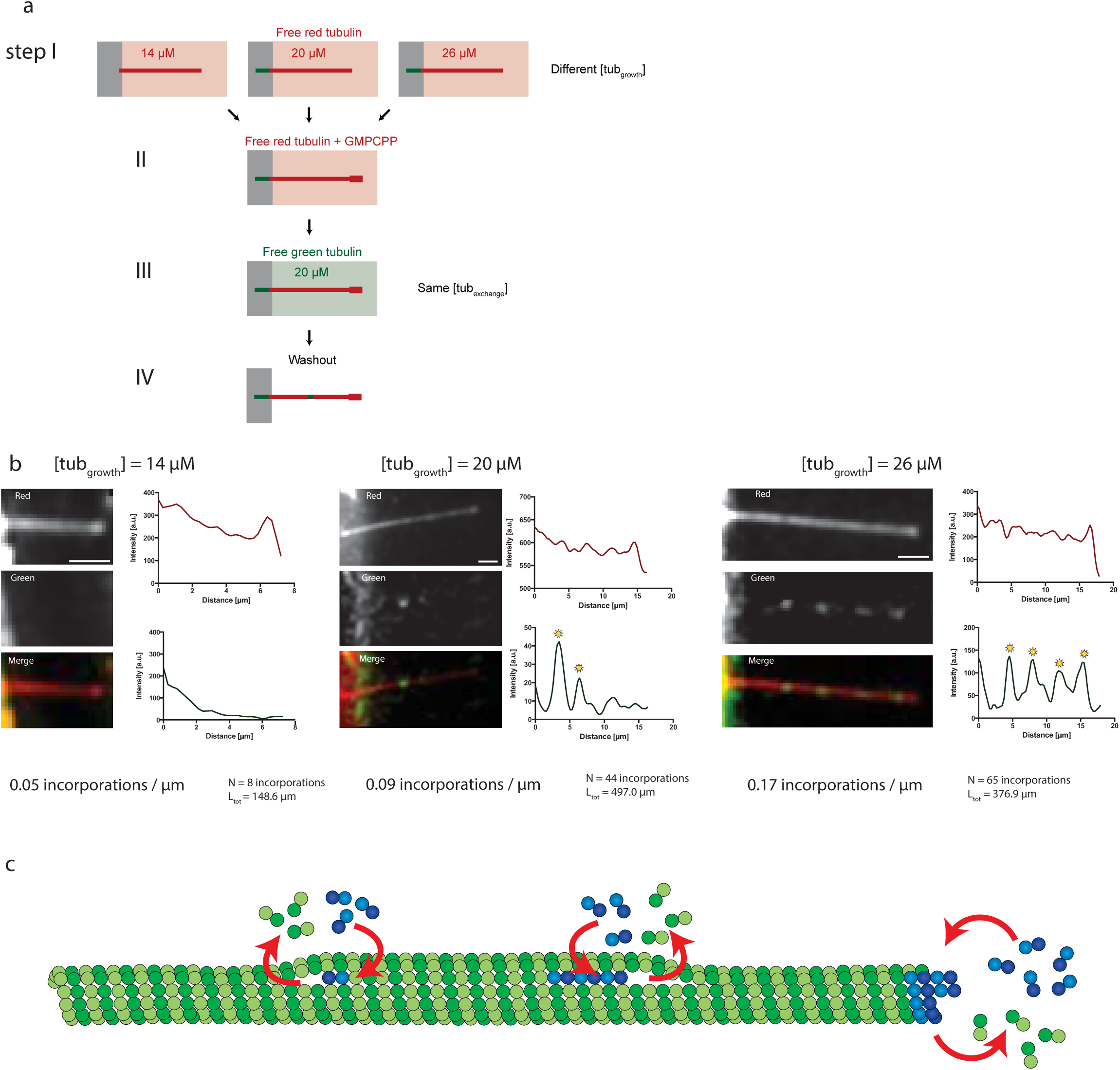
(a) The schematic representation of the experimental setup to test the role of the microtubule growth rate/lattice-defect frequency on tubulin turnover in the microtubule lattice. Microtubules were grown in 14 μM, 20 μM or 26 μM red-fluorescent free tubulin before capping with GMPCPP. Microtubules were then exposed to 20 μM green-fluorescent free tubulin for 15 min before washout. (b) The images and line scans show examples of microtubules grown at different tubulin concentrations and the average frequency of sites of tubulin incorporation. Scale bars: 3 μm (c) Model scheme for lattice dynamics at the microtubule ends and at sites of lattice defects, such as sites of transition of protofilament number.

## DISCUSSION

Taken together, our results revealed that, in a similar way to the microtubule end, the microtubule shaft can spontaneously incorporate and shed dimers (Fig 4c). Such incorporations have been observed in living cells and attributed to the effect of mechanical stress on the lattice (Aumeier et al., 2016; de Forges et al., 2016). Our results now suggest that they may occur spontaneously if the microtubule exists long enough, ie more than a dozen of minutes, which is the case in most cells in interphase and notably differentiated cells in which microtubules can exist for hours (Gelfand and Bershadsky, 1991; Li and Black, 1996). Our results also showed that turnover depends on the presence of lattice defects, which remain difficult technically to identify in living cells. However, lattice defects may be more frequent in vivo than in vitro given that the growth rate of microtubules is typically 10-fold higher in vivo (Zanic et al., 2013). On the other hand, the frequency of lattice defects in vivo may be reduced by other proteins that support tubulin assembly at the microtubule end (Akhmanova and Steinmetz, 2008), which could eliminate defects and enforce a homogeneous lattice. Nevertheless, lattice defects occur in vivo and have been shown to impact the motion of microtubule motors (Gramlich et al., 2017; Liang et al., 2016), microtubule-severing enzymes (Díaz-Valencia et al., 2011; Davis et al., 2002), and the recruitment of microtubule-associated proteins (de Forges et al., 2016). Our results suggest that dimer turnover promotes the remodeling of the microtubule's conformation around sites of lattice defects and thereby confers on it an unexpected degree of structural plasticity. Furthermore, dimer turnover, by promoting the incorporation of new GTP-bound tubulin along the lattice, is likely to promote the recruitment of specific proteins with affinity for this tubulin state and thereby potentially impact also the biochemical regulation of the microtubule network.

## Acknowledgements

This work was supported by the French National Agency for Research (ANR-16-CE11-0017-01 to DC, ANR-12-BSV5-0004-01 to MT and ANR-14-CE09-0014-02 to LB) the Human Frontier in Science Program (RGY0088 to MT) and the European Research Council (Starting Grant 310472 to MT).

## Author Contributions

LS performed all dimer exchange and fracture experiments with the help of JG. LS, LB and MT designed these experiments. DC designed and performed cryo-electron microscopy experiments. KJ designed and performed numerical simulations. LS, CA, LB, MT and KJ analyzed data. LS, MT and KJ wrote the manuscript.

## Materials and Methods

### Tubulin purification and labeling

Tubulin was purified from fresh bovine brain by three cycles of temperature-dependent assembly and disassembly in Brinkley Buffer 80 (BRB80 buffer; 80 mM PIPES, pH 6.8, 1 mM EGTA, and 1 mM MgCl_2_ plus 1 mM GTP; Shelanski, 1973). MAP-free neurotubulin was purified by cation-exchange chromatography (EMD SO, 650 M, Merck) in 50 mM PIPES, pH 6.8, supplemented with 1 mM MgCl_2_, and 1 mM EGTA (Malekzadeh-Hemmad et al., 1993). Purified tubulin was obtained after a cycle of polymerization and depolymerization. Fluorescent tubulin (ATTO-488 and ATTO-565-labeled tubulin) and biotinylated tubulin were prepared as previously described (Hyman et al., 1991). Microtubules from neurotubulin were polymerized at 37°C for 30 min and layered onto cushions of 0.1 M NaHEPES, pH 8.6, 1 mM MgCl_2_, 1 mM EGTA, 60% v/v glycerol, and sedimented by high centrifugation at 30°C. Then microtubules were resuspended in 0.1 M NaHEPES, pH 8.6, 1 mM MgCl_2_, 1 mM EGTA, 40% v/v glycerol and labeled by adding 1/10 volume 100 mM NHS-ATTO (ATTO Tec), or NHS-Biotin (Pierce) for 10 min at 37°C. The labeling reaction was stopped using 2 volumes of 2x BRB80, containing 100 mM potassium glutamate and 40% v/v glycerol, and then microtubules were sedimented onto cushions of BRB80 supplemented with 60% glycerol. Microtubules were resuspended in cold BRB80. Microtubules were then depolymerised and a second cycle of polymerization and depolymerization was performed before use.

### Cover-glass micropatterning

The micropatterning technique was adapted from Portran et al. (2013). Cover glasses were cleaned by successive chemical treatments: 30 min in acetone, 15 min in ethanol (96%), rinsing in ultrapure water, 2 h in Hellmanex III (2% in water, Hellmanex), and rinsing in ultrapure water. Cover glasses were dried using nitrogen gas flow and incubated for three days in a solution of tri-ethoxy-silane-PEG (30 kDa, PSB-2014, creative PEG works) 1 mg/ml in ethanol 96% and 0.02% of HCl, with gentle agitation at room temperature. Cover glasses were then successively washed in ethanol and ultrapure water, dried with nitrogen gas, and stored at 4°C. Passivated cover glasses were placed into contact with a photomask (Toppan) with a custom-made vacuum-compatible holder and exposed to deep UV (7 mW/cm^2^ at 184 nm, Jelight) for 2 min 30 s. Deep UV exposure through the transparent micropatterns on the photomask created oxidized micropatterned regions on the PEG-coated cover glasses.

### Microfluidic circuit fabrication and flow control

The microfluidic device was fabricated in PDMS (Sylgard 184, Dow Corning) using standard photolithography and soft lithography. The master mold was fabricated by patterning 50-μm thick negative photoresist (SU8 2100, Microchem, MA) by photolithography (Duffy et al., 1998). A positive replica was fabricated by replica molding PDMS against the master. Prior to molding, the master mold was silanized (trichloro(1H,1H,2H,2H-perfluorooctyl)silane, Sigma) for easier lift-off. Four inlet and outlet ports were made in the PDMS device using 0.5 mm soft substrate punches (UniCore 0.5, Ted Pella, Redding, CA). Connectors to support the tubing were made out of PDMS cubes (0.5 cm side length) with a 1.2 mm diameter through hole. The connectors were bonded to the chip ports using still liquid PDMS as glue, which was used to coat the interface between the chip and the connectors, and was then rapidly cured on a hotplate at 120°C. Teflon tubing (Tefzel, inner diameter: 0.03″, outer diameter: 1/16″, Upchurch Scientific) was inserted into the two ports serving as outlets. Tubing with 0.01″ inner and 1/16″ outer diameter was used to connect the inlets via two three-way valves (Omnifit labware, Cambridge, UK) that could be opened and closed by hand to a computer-controlled microfluidic pump (MFCS-4C, Fluigent, Villejuif, France). Flow inside the chip was controlled using the MFCS-Flex control software (Fluigent). Custom rubber pieces that fit onto the tubing were used to close the open ends of the outlet tubing when needed.

### Microtubule growth on micropatterns

Microtubule seeds were prepared at 10 μM tubulin concentration (30% ATTO-488-labeled or ATTO-565-labeled tubulin and 70 % biotinylated tubulin) in BRB80 supplemented with 0.5 mM GMP-CPP at 37°C for 1 h. The seeds were incubated with 1 μM Taxotere (Sigma) at room temperature for 30 min and were then sedimented by high centrifugation at 30°C and resuspended in BRB80 supplemented with 0.5 mM GMP-CPP and 1 μM Taxotere. Seeds were stored in liquid nitrogen and quickly warmed to 37°C before use.

The PDMS chip was placed on a micropatterned cover glass and fixed on the microscope stage. The chip was perfused with neutravidin (25 μg/ml in BRB80; Pierce), then washed with BRB80, passivated for 20 s with PLL-g-PEG (Pll 20K-G35-PEG2K, Jenkam Technology) at 0.1 mg/ml in 10 mM Na-Hepes (pH = 7.4), and washed again with BRB80. Microtubule seeds were flowed into the chamber at high flow rates perpendicularly to the micropatterned lines to ensure proper orientation of the seeds. Non-attached seeds were washed out immediately using BRB80 supplemented with 1% BSA. Seeds were elongated with a mix containing 14, 20 or 26 μM of tubulin (30% labeled) in BRB80 supplemented with 50 mM NaCl, 25 mM NaPi, 1 mM GTP, an oxygen scavenger cocktail (20 mM DTT, 1.2 mg/ml glucose, 8 μg/ml catalase and 40 μg/ml glucose oxidase), 0.1% BSA and 0.025% methyl cellulose (1500 cp, Sigma). GMPCPP caps were grown by supplementing the before-mentioned buffer for with 0.5 mM GMPCPP (Jena Bioscience) and using 10 μM tubulin (100% labeled with a red fluorophore). For fracture experiments, this buffer was then replaced by a buffer containing the same supplements, but without free tubulin and GTP (“washing buffer”). For incorporation experiments, the same buffer as for seed elongation was used, using 7 μM, 14 μM or 20 μM tubulin (100% labelled, green fluorescent). Microtubules were incubated in this buffer for 15 min before replacing it with washing buffer for imaging.

### Imaging

Microtubules were visualized using an objective-based azimuthal ilas2 TIRF microscope (Nikon Eclipse Ti, modified by Roper Scientific) and an Evolve 512 camera (Photometrics). The microscope stage was kept at 37°C using a warm stage controller (LINKAM MC60). Excitation was achieved using 491 and 561 nm lasers (Optical Insights). Time-lapse recording was performed using Metamorph software (version 7.7.5, Universal Imaging). Movies were processed to improve the signal/noise ratio (smooth and subtract background functions of ImageJ, version 1.47n5). To visualize incorporation, images were typically taken every 150 ms and 30 images were overlain and averaged. For fracture experiments, images were taken every 10 s.

### Cryo-electron microscopy

Analysis of lattice-defect frequency as a function of tubulin concentration was performed on microtubules assembled from phosphocellulose-purified bovine-brain tubulin nucleated by centrosomes isolated from KE-37 human lymphoid cells (Chrétien *et al*. 1995). Briefly, centrosomes and tubulin at the desired concentration (6.5 μM, 16 μM and 19 μM) were mixed and incubated directly on the EM grid under controlled humidity and temperature conditions (Chrétien *et al.*, 1992), blotted with a filter paper to form a thin film of suspension, and quickly plunged into liquid ethane. Microtubules were visualized with a EM 400 electron microscope (Philips) operating at 80 kV. Images were taken at ~1.5 μm underfocus on negatives (SO-163, Kodak). Sites of transition of protofilament and/or helix-start numbers were determined on printed views of the negatives (Chrétien *et al.*, 1992; Chrétien and Fuller, 2000).

Cryo-electron tomography was performed on microtubules self-assembled from phosphocellulose-purified porcine brain tubulin (Weiss *et al.*, 2010). Ten nm gold nanoparticles coated with BSA (Aurion) were added to the suspension to serve as fiducial markers (Coquelle *et al.*, 2011). Specimens in 4 μm aliquots were pipetted at specific assembly times and vitrified as described above. Specimen grids were observed with a Tecnai G^2^ T20 Sphera (FEI) operating at 200 kV. Tilt series, typically in the angular range ±60°, were acquired in low electron-dose conditions using a 2k × 2k CCD camera (USC1000, Gatan). Three-dimensional reconstructions were performed using the eTomo graphical user interface of the IMOD software package (Mastronarde, 1997).

### Monte-Carlo Simulations

Kinetic Monte-Carlo simulations were performed using a rejection-free random-selection method (Lukkien et al., 1998).

#### Lattice Structure

We used the same lattice structure as van Buren et al. (vanBuren et al., 2002) or Gardner et al. (Gardner et al., 2011). Briefly, the microtubule is modeled on the scale of the tubulin dimer as the canonical 13-3 (13 protofilament, 3 start left-handed helix) structure. In addition, we postulated the existence of a less stable 12-3 (12 protofilament, 3 start helix) (Chrétien and Wade, 1991; Sui and Downing, 2010). The 13-3 lattice and the 12-3 lattice were connected via dislocation defects, where a single protofilament was lost or added (Chrétien et al., 1992). All lattice structures were modeled as square lattices, i.e. each dimer has two longitudinal and two lateral neighbors. Individual lattice sites on the square lattice were identified by a doublet of integers (i,j). The lattice was periodic in a direction perpendicular to the long axis of the microtubule with an offset of 3/2 lattice sites to reproduce the seam structure. Therefore lattice sites at the seam have 2 nearest “half” neighbors across the seam, i.e. dimers at the left seam in Fig. 2a with the doublet (1,j) are in contact with dimers (13,J+2) and (13,J+1) and dimers at the right seam with doublet (13J) are in contact with dimers (1J-1) and(1,j-2) for a 13 protofilament lattice. At a dislocation defect a lattice site has only one longitudinal neighbor site.

#### Lattice transitions and rate constants

Lattice sites can be either empty or occupied by GTP-bound (T) or GDP-bound (D) dimers. Dimers interact with other dimers on nearest neighbor lattice sites via attractive interactions, characterized by bond energies *ΔG*_*1*_ and *ΔG*_*2*_ for longitudinal and lateral bonds, respectively.

We assumed that longitudinal bonds of T-T contacts were further stabilized by the energy, *ΔG*_*TT*_, following the allosteric model proposed by Alushin et al. (Alushin et al., 2014) and that independent of the occupation of the neighbor lattice site, GDP-dimers in the kinked deformation are less stable in the lattice compared with GTP dimers by the (small) energy, *ΔG*_*D*_ (Alushin et al. 2014). Furthermore, we assumed a small destabilizing energy,*ΔG*_*C*_*i*__, for dimers in the 12 protofilament lattice compared with the 13 protofilament lattice, and a destabilizing energy, *ΔG*_*S*_*i*__, for the dimer located directly at the dislocation defect.

For the passive process of polymerization and depolymerization, the principle of detailed balance was assumed to hold and therefore on and off rate constants were coupled by the relation (De Groot and Mazur, 1984)

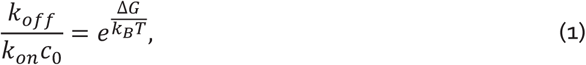

where *c*_*0*_ denotes the standard concentration of free tubulin in solution, ΔG denotes the change in free energy upon transferring a free dimer from the solution into the lattice. Δ*G* = Δ*G*_*B*_ + Δ*G*_*E*_+& contained contributions from binding of the dimer to nearest neighbors Δ*G*_*B*_*I*__, the loss of entropy due to immobilization of the free dimer in the lattice Δ*G*_*E*_*I*__, and further contributions related to the conformation of the lattice, Δ*G*_*C*_*I*__, and the presence of defects Δ*G*_*S*_. For practical reasons, we rewrote equation (1) into

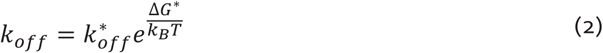

with 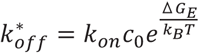 and where Δ *G*^*^ contains now only binding and conformational contributions. For the off-rate constant of a GDP dimer we assumed

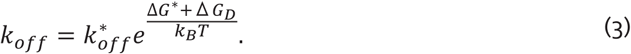

And we assumed GTP dimers are irreversibly hydrolyzed into GDP dimers by the rate constant *k*_*hy*_ if their hydrolysable β subunit is in contact with the α subunit of another dimer (i.e. if the top-next longitudinal lattice site is occupied). Hence our modelling permitted the following transitions: GTP dimers that can polymerize into a lattice structure and depolymerize or hydrolyze into GDP dimers; and GDP dimers that can depolymerize from the lattice.

At dislocation defects, the lattice can breathe (i.e. open with rate constant k_br_ to create an empty lattice site and close with rate constant k_cl_ to annihilate an empty lattice site) as shown in Fig. 2e. Because breathing is a reversible process k_br_ and k_cl_ are related by

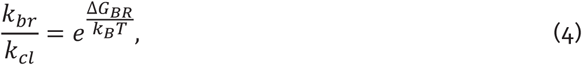

where *ΔG*_*BR*_ denotes the difference in lattice free energy between the closed and open lattice state and it contains contributions from the conformational energy penalty for a 12 protofilament lattice, which accelerates opening because a less favorable 12 protofilament lattice is receding in favor of a more stable 13 protofilament lattice, and contributions from the lateral binding energy of the bond which will be opened and the release in defect strain energy. *ΔG*_*BR*_ depends on the occupation of the lattice sites in proximity to the defect.

Monte-Carlo simulations were performed using a custom-written C code. Unless stated otherwise, we used the following parameters listed in Table S1.

**Table S1:** Parameter values used for the Monte-Carlo simulations unless stated otherwise.

| Parameter | Value | References/Remarks |
| --- | --- | --- |
| $\Delta G_1$ | $-14.4 k_B T$ | $\Delta G_1 + \Delta G_{TT} + \Delta G_E = -8.8 k_B T$ (Gardner et al., 2011) |
| $\Delta G_{TT}$ | $-5.6 k_B T$ | adapted for our model (van Buren et al., 2002) |
| $\Delta G_2$ | $-5 k_B T$ | Gardner et al., 2011 |
| $\Delta G_E$ | $11.2 k_B T$ | lowest estimate is $10 k_B T$ (Howard et al., 2001, vanBuren et al., 2002, Erickson, 1989) |
| $\Delta G_D$ | $1 k_B T$ | adapted for our model |
| $\Delta G_C$ | $0.2 k_B T$ | adapted for our model (Hunyadi et al. , 2005) |
| $\Delta G_S$ | $0-3.75 k_B T$ | adapted for our model |
| $k_{on}$ | $0.4 \mu M^{-1} s^{-1}$ | adapted for our model to give an approximate microtubule elongation rate of $2 \mu m \cdot min^{-1}$, overall on-rate per 13_3 microtubule is $k_{on}^{MT} = 5.2 \mu M^{-1} s^{-1}$ (Gardner et al. 2011, vanBuren et al. 2002, Melki et al. 1996, Walker et al. 1988) |
| $k_{off}^*$ | $2.86 \times 10^{10} s^{-1}$ | |
| $k_{off}$ | $0.4 s^{-1}$ | for a GTP dimer with one lateral and one longitudinal GTP-bound neighbor [for a GTP dimer with one longitudinal GTP-bound neighbor $k_{off} = 60 s^{-1}$] |
| $k_{hy}$ | $0.4 s^{-1}$ | Melki et al. 1996 |
| $k_{cl}$ | $0.7 s^{-1}$ | fully occupied lattice at the dislocation |
| $k_{br}$ | $0.4 s^{-1}$ | fully occupied lattice at the dislocation |

## REFERENCES

Akhmanova, A., and M.O. Steinmetz. 2008. Tracking the ends: a dynamic protein network controls the fate of microtubule tips. Nat. Rev. Mol. Cell Biol. 9:309–322. doi:10.1038/nrm2369.

Aumeier, C., L. Schaedel, J. Gaillard, K. John, L. Blanchoin, and M. Théry. 2016. Self-repair promotes microtubule rescue. Nat. Cell Biol. 18:1054–1064. doi:10.1038/ncb3406.

Chrétien, D., and R.H. Wade. 1991. New data on the microtubule surface lattice. Biol. Cell. 71: 161–174.

Chrétien, D., and S.D. Fuller. 2000. Microtubules switch occasionally into unfavorable configurations during elongation. J. Mol. Biol. 298:663–76. doi:10.1006/jmbi.2000.3696.

Chrétien, D., F. Metoz, F. Verde, E. Karsenti, and R.H. Wade. 1992. Lattice defects in microtubules: protofilament numbers vary within individual microtubules. J. Cell Biol. 117:1031–40.

Chrétien, D., S.D. Fuller, and E. Karsenti. 1995. Structure of growing microtubule ends: two-dimensional sheets close into tubes at variable rates. J. Cell Biol. 129: 1311–1328.

Coquelle F., Blestel S., Heichette C., Arnal I., Kervrann C. and D. Chrétien. 2011. Cryo-electron tomography of microtubules assembled in vitro from purified components. Methods in Molecular Biology. 777:193–208. doi:10.1007/978-1-61779-252-64.

Davis, L.J., D.J. Odde, S.M. Block, and S.P. Gross. 2002. The Importance of Lattice Defects in Katanin-Mediated Microtubule Severing in Vitro. Biophys. J. 82:2916–2927.

Diaz-Valencia, J.D., M.M. Morelli, M. Bailey, D. Zhang, D.J. Sharp, and J.L. Ross. 2011. Drosophila katanin-60 depolymerizes and severs at microtubule defects. Biophys. J. 100:2440–9. doi:10.1016/j.bpj.2011.03.062.

Doodhi, H., A.E. Prota, R. Rodriguez-Garcia, H. Xiao, D.W. Custar, K. Bargsten, E.A. Katrukha, M. Hilbert, S. Hua, K. Jiang, I. Grigoriev, C.-P.H. Yang, D. Cox, S.B. Horwitz, L.C. Kapitein, A. Akhmanova, and M.O. Steinmetz. 2016. Termination of Protofilament Elongation by Eribulin Induces Lattice Defects that Promote Microtubule Catastrophes. Curr. Biol. 26:1713–1721. doi:10.1016/j.cub.2016.04.053.

Duellberg, C., N.I. Cade, D. Holmes, and T. Surrey. 2016. The size of the EB cap determines instantaneous microtubule stability. Elife. 5:1–23. doi:10.7554/eLife.13470.

Dye, R.B., P.F. Flicker, D.Y. Lien, and R.C. Williams. 1992. End-stabilized microtubules observed in vitro: stability, subunit, interchange, and breakage. Cell Motil. Cytoskeleton. 21:171–86. doi:10.1002/cm.970210302.

de Forges, H., A. Pilon, I. Cantaloube, A. Pallandre, A.-M. Haghiri-Gosnet, F. Perez, and C. Poüs. 2016. Localized Mechanical Stress Promotes Microtubule Rescue. Curr. Biol. 26:3399–3406. doi:10.1016/j.cub.2016.10.048.

Gardner, M.K., B.D. Charlebois, I.M. Janosi, J. Howard, A.J. Hunt, and D.J. Odde. 2011. Rapid microtubule self-assembly kinetics. Cell 146: 582–592.

Gelfand, V.I., and A.D. Bershadsky. 1991. Microtubule Dynamics: Mechanism, Regulation, and Function. Annu. Rev. Cell Biol. 7:93–116. doi:10.1146/annurev.cb.07.110191.000521.

Gramlich, M.W., L. Conway, W.H. Liang, J.A. Labastide, S.J. King, J. Xu, and J.L. Ross. 2017. Single Molecule Investigation of Kinesin-1 Motility Using Engineered Microtubule Defects. Sci. Rep. 7:44290. doi:10.1038/srep44290.

Guesdon A., Bazile F., Buey R.M., Mohan R., Monier S., Rodriguez Garcia R., Angevin M., Heichette C., Wieneke R., Tampé R., Duchesne L., Akhmanova A., Steinmetz M.O., and D. Chrétien. 2016. EB1 interacts with outwardly curved and straight regions of the microtubule lattice. Nature Cell Biology 18:1102–8. doi: 10.1038/ncb3412.

Hunyadi, V., Chrétien, D. and I. M. Janosi. 2005. Mechanical stress induced mechanism of microtubule catastrophes. J. Mol. Biol. 348:927–938.

Inoue, S. 1981. Cell division and the mitotic spindle. J. Cell Biol. 91:131–147.

Kellogg, E.H., N.M.A. Hejab, S. Howes, P. Northcote, J.H. Miller, J.F. Diaz, K.H. Downing, and E. Nogales. 2017. Insights into the Distinct Mechanisms of Action of Taxane and Non-Taxane Microtubule Stabilizers from Cryo-EM Structures. J. Mol. Biol. 429:633–646. doi:10.1016/j.jmb.2017.01.001.

Leslie, R.J., W.M. Saxton, T.J. Mitchison, B. Neighbors, E.D. Salmon, and J.R. McIntosh. 1984. Assembly properties of fluorescein-labeled tubulin in vitro before and after fluorescence bleaching. J. Cell Biol. 99:2146–2156. doi:10.1083/jcb.99.6.2146.

Li, Y., and M.M. Black. 1996. Microtubule assembly and turnover in growing axons. J. Neurosci. 16:531–544.

Liang, W.H., Q. Li, K.M.R. Faysal, S.J. King, A. Gopinathan, and J. Xu. 2016. Article Microtubule Defects Influence Kinesin-Based Transport In Vitro. Biophys. J. 110:2229–2240. doi:10.1016/j.bpj.2016.04.029.

Mandelkow, E.-M., R. Schultheiss, R. Rapp, M. Müller, and E. Mandelkow. 1986. On the surface lattice of microtubules: helix starts, protofilament number, seam, and handedness. J. Cell Biol. 102:1067–1073.

Mitchison, T., and M.W. Kirschner. 1984. Dynamic instability of microtubule growth. Nature. 312:237–42.

Portran, D., J. Gaillard, M. Vantard, and M. Théry. 2013. Quantification of MAP and molecular motor activities on geometrically controlled microtubule networks. Cytoskeleton (Hoboken). 70:12–23. doi:10.1002/cm.21081.

Reid, T.A., C. Coombes, and M.K. Gardner. 2017. Manipulation and Quantification of Microtubule Lattice Integrity. Biology Open. 6: 1245–1256.

Salmon, E.D., R.J. Leslie, W.M. Saxton, M.L. Karow, and J.R. McIntosh. 1984. Spindle microtubule dynamics in sea urchin embryos: analysis using a fluorescein-labeled tubulin and measurements of fluorescence redistribution after laser photobleaching. J. Cell Biol. 99:2165–74.

Schaap, I.T., P.J. de Pablo, and C.F. Schmidt. 2004. Resolving the molecular structure of microtubules under physiological conditions with scanning force microscopy. Eur. Biophys. J. 33:462–7. doi:10.1007/s00249-003-0386-8.

Schaedel, L., K. John, J. Gaillard, M. V Nachury, L. Blanchoin, and M. Théry. 2015. Microtubules selfrepair in response to mechanical stress. Nat. Mater. 14:1156–1163. doi:10.1038/nmat4396.

Sept, D., N.A. Baker, and J.A. McCammon. 2009. The physical basis of microtubule structure and stability. Protein Sci. 12:2257–2261. doi:10.1110/ps.03187503.

Soltys, B.J., and G.G. Borisy. 1985. Polymerization of tubulin in vivo: Direct evidence for assembly onto microtubule ends and from centrosomes. J. Cell Biol. 100:1682–1689.

VanBuren, V., D.J. Odde, and L. Cassimeris. 2002. Estimations of lateral and longitudindal bond energies within the microtubule lattice. Proc. Natl. Acad. Sci. USA 99: 6035–6040.

VanBuren, V., L. Cassimeris, and D.J. Odde. 2005. Mechanochemical model of microtubule structure and self-assembly kinetics. Biophys. J. 89:2911–2926. doi:10.1529/biophysj.105.060913.

Vitre, B., F.M. Coquelle, C. Heichette, C. Garnier, D. Chrétien, and I. Arnal. 2008. EB1 regulates microtubule dynamics and tubulin sheet closure in vitro. Nat. Cell Biol. 10:415–21. doi:10.1038/ncb1703.

Weisenberg, R.C. 1972. Microtubule Formation in vitro in Solutions Containing Low Calcium Concentrations. Source Sci. New Ser. 177174254:1104–1105.

Wu, Z., H.-H. Wang, W. PLoS ONE Mu, Z. Ouyang, E. Nogales, and J. Xing. 2009. Simulations of tubulin sheet polymers as possible structural intermediates in microtubule assembly. 4 (10): e7291.

Yajima, H., T. Ogura, R. Nitta, Y. Okada, C. Sato, and N. Hirokawa. 2012. Conformational changes in tubulin in GMPCPP and GDP-taxol microtubules observed by cryoelectron microscopy. J. Cell Biol. 198:315–322. doi:10.1083/jcb.201201161.

Zanic, M., P.O. Widlund, A. a Hyman, and J. Howard. 2013. Synergy between XMAP215 and EB1 increases microtubule growth rates to physiological levels. Nat. Cell Biol. 15:688–93. doi:10.1038/ncb2744.

## References

De Groot, S.R., and P. Mazur. 1984. Non-equilibrium thermodynamics. Dover Publications Inc., New York.

Duffy, D.C., J.C. McDonald, O.J. Schueller, and G.M. Whitesides. 1998. Rapid prototyping of microfluidic systems in poly(dimethylsiloxane). Anal. Chem. 70: 4974–4984.

Erickson, H.P. 1989. Co-operativity in protein-protein association: The structure and stability of the actin filament. J. Mol. Biol. 206: 465–474.

Howard, J. 2001. Mechanics of motor proteins and the cytoskeleton. Sinauer Associates Sunderland, MA.

Hyman, A., D. Drechsel, D. Kellogg, S. Salser, K. Sawin, P. Steffen, L. Wordeman, and T. Mitchison. 1991. Preparation of modified tubulins. Methods Enzymol. 196: 478–485.

Mastronarde, D.N. 1997. Dual-axis tomography: an approach with alignment methods that preserve resolution. J. Struct. Biol. 120: 343–352.

Lukkien, J.J., J.P.L. Segers, P.A.J. Hilbers, R.J. Gelten, and A.P.J. Jansen. 1998. Efficient Monte Carlo methods for the simulation of catalytic surface reactions. Phys. Rev. E 58: 2598–2610.

Malekzadeh-Hemmat, K., P. Gendry, and J.F. Launey. 1993. Rat pankreas kinesin: Identification and potential binding to microtubules. Cell. Mol. Biol. 39: 279–285.

Melki, R., S. Fievez, and M.-F. Carlier. 1996. Continuous monitoring of Pi-release following nucleotide hydrolysis in actin or tubulin assembly using 2-amino-6-mercapto-7-methylpurine ribonucleoside and purine-nucleoside phosphorylase as an enzyme linked assay. Biochem. 35: 12038–12045.

Pettersen E.F., T.D. Goddard, C.C. Huang, G.S. Couch, D.M. Greenblatt, E.C. Meng, and T.E. Ferrin. 2004. CSF Chimera‐‐a visualization system for exploratory research and analysis. J. Comput. Chem. 25:1605–12.

Shelanski, M.L. 1973. Chemistry of the filaments and tubules of brain. J. Histochem. Cytochem. 21: 529–539.

Walker, R.A., E.T. O'Brien, N.K. Pryer, M.F. Soboeiro, W.A. Voter, H.P. Erickson, and E.D. Salmon. 2002. Dynamic instability of individual microtubules analyzed by video lighmicroscopy: Rate constants and transition frequencies. J. Cell Biol. 107: 1437–1448.

Weis, F., L. Moullintraffort, C. Heichette, D. Chrétien, and C. Garnier. 2010. The 90-kDa heat shock protein HSP90 protects tubulin against thermal denaturation. J. Biol. Chem. 285:9525–34.

